# Markov state models from hierarchical density-based assignment

**DOI:** 10.1101/2021.05.13.444064

**Authors:** Ion Mitxelena, Xabier López, David de Sancho

## Abstract

Markov state models (MSMs) have become one of the preferred methods for the analysis and interpretation of molecular dynamics (MD) simulations of conformational transitions in biopolymers. While there is great variation in terms of implementation, a well-defined workflow involving multiple steps is often adopted. Typically, molecular coordinates are first subjected to dimensionality reduction and then clustered into small “microstates”, which are subsequently lumped into “macrostates” using the information from the slowest eigenmodes. However, the microstate dynamics is often non-Markovian and long lag times are required to converge the MSM. Here we propose a variation on this typical workflow, taking advantage of hierarchical density-based clustering. When applied to simulation data, this type of clustering separates high population regions of conformational space from others that are rarely visited. In this way, density-based clustering naturally implements assignment of the data based on transitions between metastable states. As a result, the state definition becomes more consistent with the assumption of Markovianity and the timescales of the slow dynamics of the system are recovered more effectively. We present results of this simplified workflow for a model potential and MD simulations of the alanine dipeptide and the FiP35 WW domain.

## I. INTRODUCTION

Atomistic molecular dynamics (MD) simulations of biomolecular systems are part of the essential toolbox of modern biophysics^1^. Traditionally, this technique has faced multiple challenges in terms of accuracy and precision, due to problems in energy functions (“force fields”) and the short integration time-step required in the simulations, respectively. However, in the last decades we have witnessed enormous progress in both fronts^2–5^. These improvements leave us with the additional burden of having to analyze extremely large data sets including Cartesian coordinates of many-atom systems with (potentially) femtosecond resolution. In the last 15 years, Markov State Models (MSMs) have become established as a useful approach for both the analysis and interpretation of MD simulations^6–8^. Starting from one or multiple MD trajectories, MSMs provide an integrated framework for constructing a network model of metastable states such that biomolecular dynamics are described as memory-less jump processes. From the network model, equilibrium and dynamic properties are accessible, which allows comparison against experiment. Using transition path theory^6^, one can recover an intuitive grasp of the mechanism of the rare events of interest. Today, a variety of software packages are available for the construction of MSMs^9–11^.

Although Markov state modelling is an area of very active development, typically, the construction of an MSM involves a well-defined number of steps^12,13^. These include featurization, dimensionality reduction, clustering and discretization into fine grained microstates, estimation and, finally, coarsegraining into intuitively understandable models. Each of these steps require decisions from the practicioner. One potential problem in this protocol is that the microstates will often not fulfil the Markovian assumption that the methodology builds upon, as they do not match true metastable basins where the molecule loses memory before visiting the next microstate. This issue was addressed by Noé and co-workers in the construction of hidden Markov models and observable operator models^14,15^, and by Guarenera and Vanden Eijnden in the context of milestoning^16^. Furthermore, any MSM is affected of a systematic error due to the discretization of the state space^12^. Prinz et al. concluded that this discretization error can be diminished by choosing a fine discretization and using a long lag time *τ* (see below). However, they also pointed out that if only the slowest dynamical processes are of interest, it is sufficient to discretize the state space such that the first few eigenvectors are well represented.

In this work, we propose an alternative approach to these problems in the event that the fast intra-well dynamics are not important. We combine the information from optimal order parameters, which may be derived from dimensionality reduction for molecular systems, and hierarchical density-based clustering methods to directly define states that better fulfil the assumption of Markovianity. In this way, we obtain in a single step coarse-grained MSMs that retain the accuracy of fine grained models. Our proposed workflow, that we term “hdbMSM”, hence compresses some of the steps defined above, hopefully making the construction of the MSM more straightforward.

The main reason for the usefulness of hdbMSMs is that the hierarchical density-based clustering implements the “transition based assignment” (TBA) introduced by Buchete and Hummer^17^. In their study of short peptide dynamics, master equation models were constructed using a discretization in torsion angle space. Buchete and Hummer defined circular high density regions corresponding to the *α*-helix and PPII/*β* conformations in the Ramachandran map to define microstates. In this way, the state assignment eliminated fast non-Markovian recrossings, resulting in drastically improved relaxation times that were independent of the lag time used in the construction of the MSM^17^. This idea is also referred to as “coring”, and has been used in the literature to maximize the metastability of discrete states^18^. In our implementation, the definition of cores is streamlined by the combination of optimal coordinates, which may come from dimensionality reduction in complex systems, and density-based clustering.

Density-based clustering has been introduced before in the literature of MSM development by Lemke and Keller^19^ and Sittel and Stock^20^. Specifically, Lemke and Keller showed that combining density-based clustering using the DBSCAN, Jarvis-Patrick and CNN algorithms, with the core-set approach reduces the approximation error^12^ and improves the spatial resolution of the models^19,21^. Similarly, Sittel and Stock developed a density-based geometrical clustering algorithm based on local free energy estimates, so that resulting microstates are separated by local free energy barriers. They also described a complete workflow to robustly generate MSMs, but the latter required as many steps as the standard approaches. More recently, Nagel and Stock exploited density-based clustering methods to develop dynamical coring of MSMs^22^. Dynamical coring aims to avoid counting intrastate fluctuations as transitions between different metastable states. For high dimensional systems, Nagel and Stock introduce an additional parameter to properly define metastable conformational states. Thus, they impose that a transition is only considered if the trajectory spends a minimum time *τ*_cor_ in the new state or core region.

Here, we exploit the HDBSCAN clustering algorithm recently developed by Campello, Moulavi and Sander^23,24^. Contrary to its predecessor, DBSCAN^25^, or the other methods used by Sittel and Stock^20^ and Lemke and Keller^19,21^, HDB-SCAN does not require a neighborhood parameter, which can be difficult to determine depending on the featurization. Instead, HDBSCAN requires a single parameter which defines the smallest size grouping that the user wishes to consider a cluster. We present our modified workflow for constructing MSMs and apply it to a number of typical problems of increasing complexity. Depending on the complexity of the system we can either apply HDBSCAN to the natural coordinates (e.g. the *ϕ* and *ψ* torsions of the alanine dipeptide) or to coordinates optimally derived using dimensionality reduction methods like TICA^26–28^. In each of these examples, we compare our results with those from a standard MSM.

## II. THEORY

In the present section we summarize only a few of the essential concepts associated to the construction of MSMs that are necessary for this paper. A more detailed discussion can be found elsewhere^6,12^.

Markovian models describe the dynamics of a set of disjoint discrete states based on a transition probability matrix

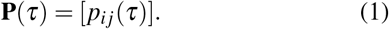

Each of the elements *p*_*i j*_(*τ*) in this matrix is the conditional probability that the system will be in state **x** _*j*_ at time *t* + *τ* if the state at time *t* is **x**_*i*_. The condition of Markovianity implies that this probability only depends on the current state and not of the rest of the trajectory of the system. In order to calculate the matrix **P**(*τ*) only a discrete state trajectory and a chosen lag time *τ* are required. Multiple methods can be found in the literature to estimate the transition probability matrix from the number of observed jumps between states^9^.

Since the conditional probabilities are explicitly dependent on *τ*, this parameter is crucial to build an MSM. The accuracy of MSMs, and therefore their validity, is often first judged by comparing the implied timescales (*t*_*i*_ or ITS)

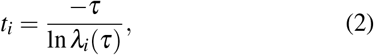

at different values of *τ*. In this expression, *λ*_*i*_(*τ*) denotes the *i*-th eigenvalue of the matrix **P**(*τ*). Since any Markovian model systematically underestimates the timescales *t*_*i*_, an adequate lag time is determined by the smallest value of *τ* among those corresponding to the saturated *t*_*i*_ values.

An additional and more stringent requirement to have a reliable MSM is the Chapman-Kolmogorov (CK) test^12^

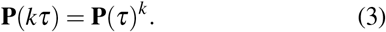

Eq. 3 allows to validate the model predictions against the observed data. Recently, the development of the variational approach for conformational dynamics (VAC)^29^ and Markov processes (VAMP)^30^ has opened up the possibility to quantitatively validate the parameters choice. Once the set of discrete states and corresponding transition probability matrix are defined, one can proceed to analyze the resulting model, e.g. by inspecting the eigenvectors and eigenvalues of **P**(*τ*). The former describe the stationary and transition modes of the system, whereas the latter are the corresponding relaxation times, as indicated by Eq. 2. Also, in the typical workflow, one will use the information in these eigenvectors to coarse-grain the MSM into an intuitively understandable model using methods like PCCA^31^.

## III. METHODS

### A. Typical and proposed workflows for MSM construction

The construction of an MSM from MD trajectories typically involves the following steps (see Fig. 1A):

**FIG. 1:**
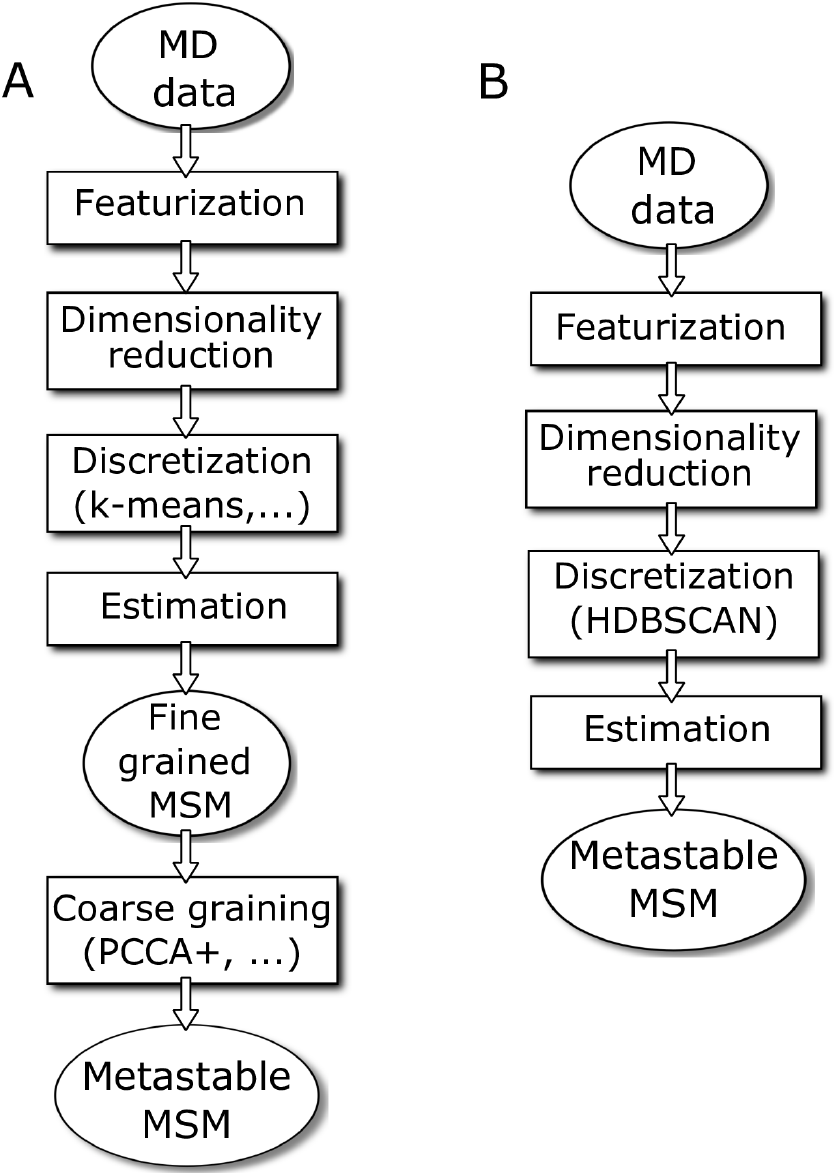
Typical (A) and proposed (B) workflows for the construction of Markov state models.

i. *Featurization*: Cartesian coordinates are transformed into more informative quantities, such as distances between heavy atoms or backbone torsion angles.
ii. *Dimensionality reduction*: A transformation of the feature data into a set of collective variables that capture most of the information. In the present work this is done by using the time-lagged independent coordinate analysis (TICA)^26–28^. Many other dimensionality reduction methods are available in the literature^7,30,32,33^.
iii. *Discretization*: The projection on the relevant coordinates is discretized using clustering methods, typically the *k*-means method^34^. The corresponding disjoint states are referred to as microstates.
iv. *Estimation*: The definition of discrete states in the previous step, together with the adoption of a proper lag time, allow us to build a transition probability matrix that describe the dynamics of the system.
v. *Coarse-graining*: In order to produce interpretable models, the MSM is reduced to a few states model by coarse-graining methods such as PCCA+^31,35^. Resulting macrostates must be associated with kinetically metastable regions.

Intimately related with the construction of an MSM is its validation. The CK test mentioned above is routinely employed to validate any given MSM, namely if its corresponding transition probability matrix is Markovian or not. Additional validation tools are the calculation of physical observables from the MSM or the comparison against reference data. Recently, the development of alternative approaches to MSMs, as the estimation of a Koopman model to compute ITS, has opened new strategies for MSM validation^30^.

In this work we present an alternative workflow for the derivation of MSMs, which compresses steps (iii) to (v) above into a single step (see Fig. 1B), by virtue of using hierarchical density-based clustering (hence the name hdbMSM). In this case, after dimensionality reduction the projections on the coordinates are clustered into a few states using hierarchical density-based clustering (see below). The resulting discrete states trajectory can be used to construct MSMs without loss in the quality of the estimation of timescales. A similar strategy has been recently adopted in the context of deep learning methods^30^.

All MSM calculations were performed using the PyEMMA Python library^9^, version 2.5.7 (http://www.emma-project.org). In all cases we report both the maximum likelihood estimate of the implied timescales and the mean from a Bayesian sampling of the posterior, from which we also derive errors with a 95% confidence interval. A collection of Python3 notebooks to reproduce our results is openly available at https://github.com/BioKT/hdbMSM.

### B. Hierarchical density-based clustering

Typically, clustering in MSM construction is performed using flat, centroid-based algorithms like *k*-means. Here we define the states in the hdbMSM using density-based clustering methods^36^. In contrast with the popular *k*-means algorithm, which performs best if the clusters are spherical and equally-sized, density-based methods are able to produce arbitrarily shaped clusters. Specifically, we use HDBSCAN (for Hierarchical Density-Based Spatial Clustering of Applications with Noise). HDBSCAN estimates a density function for each cluster and thereby clusters of different densities are allowed. In this way, HDBSCAN converts DBSCAN into a hierarchical clustering algorithm.

An advantage of HDBSCAN over other density-based methods is that it does not require the definition of non-intuitive parameters that are hard to set, as is the case of the *neighbourhood* parameter *ε*. Being hierarchical, however, HDBSCAN requires defining the level set for the hierarchy, which is done setting the *minimum cluster size*. This parameter is related to the smallest number of datapoints the user wants to consider as an independent cluster. Importantly, data that do not fulfil the requirements to be part of a cluster are classified as noise. There is an additional parameter that can be defined by the user to fine tune the clustering, the *minimum samples*. It stands for the number of samples in a neighbourhood for a point to be considered a core point. By default its value is the same as the *minimum cluster size*, nevertheless, for large datasets of tens or hundreds of thousands of entries we recommend to set it not larger than a few hundreds.

At the end of the procedure, the selected clusters are extracted from the condensed tree plot (see Results section). This plot is obtained from a conventional cluster hierarchy by saving only those clusters that are larger than the *minimum cluster size*. Then, HDBSCAN computes the stability of each cluster and keeps only the most stable ones. This is done by using *λ* = 1*/distance* to compute the persistence of each cluster, where *distance* is defined as the mutual reachability distance. So if a given cluster takes small *λ* values, it will be large in size when we represent it in the feature space. Thus, clusters look cone shaped in the condensed tree plot, as the width corresponds to the number of datapoints it contains and the size of the cluster decreases from small (top part) to larger (bottom part) *λ* values. A more detailed description of the HDBSCAN algorithm can be found elsewhere^23,24^. A Python implementation of HDBSCAN is freely available at https://hdbscan.readthedocs.io/^24^.

More importantly, HDBSCAN identifies clusters as highly probable regions separated by improbable regions, so the latter are left as noise that do not belong to any cluster. This is consistent with the spirit of the aforementioned TBA procedure for defining states in MSMs^17^. We note that HDBSCAN offers the possibility to control the amount of noise and be either more or less conservative in the assignment by computing for each point in the feature space the probability of being in its cluster. This is intimately related with soft clustering, and it is extremely useful to deal with not well-defined metastable regions, which are frequent in the conformational space of biomolecular systems.

### C. Application examples

We produce MSMs from simulation trajectories for three systems with different levels of complexity. We start from a model potential on two dimensions, then move onto the simplest peptide system undergoing molecular transitions and finally analyze a more complex dataset of long-timescale trajectories for protein folding.

#### 1. Two-dimensional potential

As a trivial model system we consider the bistable potential introduced by Berezhkovskii et al. for the study of anisotropic diffusion^37^ and later used by Cossio et al. to understand instrumental effects in single molecule force spectroscopy experiments^38^. The two dimensional potential is of the form

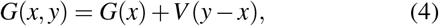

where

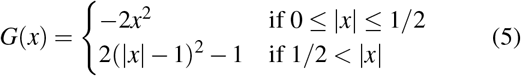

and *V* (*y* — *x*) = 0.5*κ*(*y* − *x*)^2^ where *κ* is known as the linker force constant. To generate trajectories on the two-dimensional potential described by Eq. 4, we run Brownian dynamics simulations using displacements

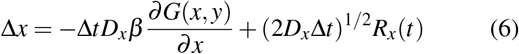

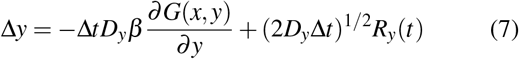

where *D*_*x/y*_ = 1 is the diffusion coefficient in the corresponding coordinate, Δ*t* is the time-step, *β* is the inverse temperature and *R*_*x/y*_(*t*) is the random force with zero mean and unit variance. The time-scales for the transitions in the model are set by *D*_*x*_Δ*t* = 2 *×* 10^−5^.

#### 2. Alanine dipeptide

We have run simulations of the terminally blocked alanine residue using the optimized Amber 99sb-ILDN force field by Lindorff-Larsen et al^39^ and the TIP3P water model^40^. The molecule was solvated in a cubic box large enough to leave 1 nm of water in each dimension. Na^+^ and Cl^−^ ions were added to approximate a 100 mM salt concentration. The system was energy minimized using a steepest descent algorithm. Then, water was equilibrated in the NVT ensemble at 300 K using the Berendsen thermostat^41^ and restraining the positions of the peptide heavy atoms. Next, the density of the box was converged at 1 Pa using the Berendsen barostat. Finally, NVT production runs were performed using a stochastic integrator and a 2 fs time step^42^. Electrostatics were calculated using the particle-mesh Ewald method^43^. The simulations were performed using the Gromacs software package^44^.

#### 3. FiP35 WW domain

We have used two independent 100 *µ*s simulation trajectories of folding/unfolding of the 32-residues long FiP35 variant of the Pin1 WW domain by Shaw and co-workers^45^. For these simulations the Amber99sb-ILDN force field^39^ and the TIP3P water model^40^ were employed. Simulations were carried out at 395 K using the Anton supercomputer. Further details are available elsewhere^45^.

## IV. RESULTS

Below we show the results obtained for the three systems introduced above using the proposed hdbMSM workflow. For reference, in all cases we first present results using the typical workflow summarized in Fig. 1A. For each of the systems complementary figures for the validation of the MSMs are included in the supplementary material.

### A. Two-dimensional potential

We start by applying the typical and proposed workflows for the construction of MSMs to a dataset derived from Brownian dynamics simulations on the double well potential by Berezhkovskii et al.^37,38^ (see Figure S1 in the Supporting Information). Since there are only two degrees of freedom, we can directly use them in the discretization. In this type of model system, one would normally assign the data points to microstates using a grid or perform *k*-means clustering. In Figure 2A we show the results from the latter approach using *k* = 100 cluster centres. From the time series data assigned to the collection of microstates, we have estimated the fine grained MSM, which we then coarse-grained using PCCA+ (i.e. following the procedure outlined in Fig. 1A). The results from the PCCA+ coarse-graining and the implied time-scales are shown in Figure 2A and D, respectively.

**FIG. 2:**
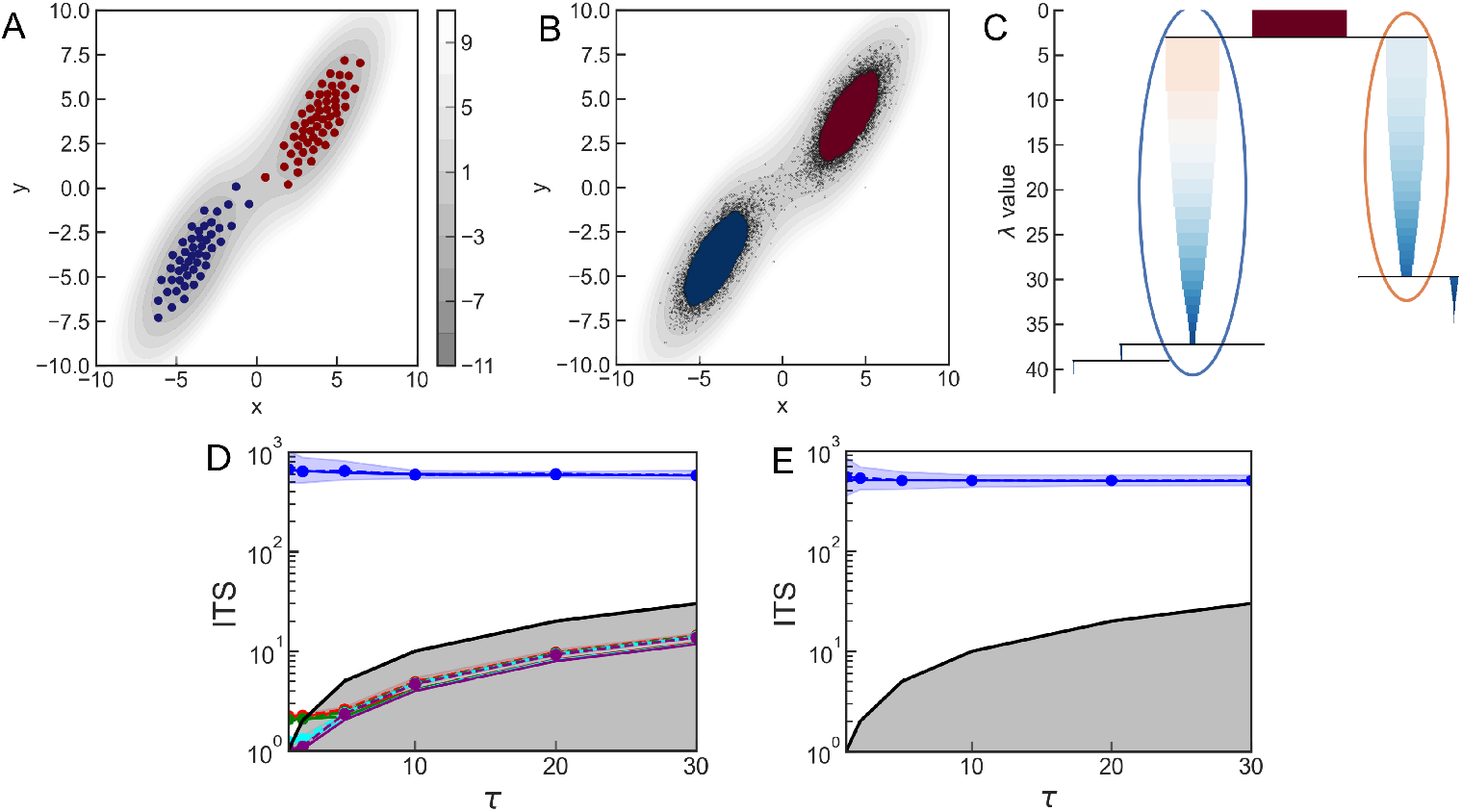
(A) Values of *x* and *y* for the 100 *k*-means clusters used in the fine grained MSM. Colours correspond to the two metastable states derived from PCCA+. The gray contour plot is the *G*(*x, y*) potential in units of *k*_*B*_*T* calculated from the analytical expressions. (B) Simulation data points coloured by the HDBSCAN assignment, using a minimum cluster size of 50. Metastable states are in red and blue and noise points are shown in black. (C) Condensed tree plot from the clustering of the data using HDBSCAN using a minimum cluster size of 50. Selected clusters are circled. (D) Implied timescales for the first five eigenmodes as a function of the lag time *τ*. The 1:1 line is shown in black for reference. (E) Same for the hdbMSM. Solid lines are maximum likelihood estimates and dashed lines are mean estimates from Bayesian sampling. Shaded areas correspond to 95% confidence intervals. The black line marks the 1:1 line, below which no reliable information can be obtained from the MSM.

Alternatively, we can simply resort to HDBSCAN to generate two core states and directly construct an accurate coarse-grained MSM. In Figure 2B we show the two core states from the hdbMSM. The assignment resulting from HDB-SCAN very much matches that from the PCCA+ coarse-graining, but leaves out as “noise” states in the barrier region and walls, directly implementing the transition based assignment method^17^. In Figure 2C we show the condensed tree plot generated from the data, which shows how the two main clusters are easily identified as those having the largest persistence within the hierarchy. In this case we further refined the amount of noise in the assignment using the probabilities that the datapoints were assigned to the clusters. Specifically, we left as noise points that had probabilities lower than 0.4.

In Figure 2D-E, we compare the implied timescales derived from both approaches. The only relevant timescale in this system, corresponding to the crossing of the barrier between both basins, is captured equally well by both the fine-grained MSM (Figure 2D) and our direct approach (Figure 2E). In both cases we find the implied timescale to be entirely independent of the lag time for the construction of the model. We have performed Chapman-Kolmogorov tests for both MSMs (see Supplementary Information, Fig. S2 and S3). In both cases Eq. 3 is satisfied accurately, as we recover excellent agreement between estimation and prediction curves. It is worth noting that when using hdbMSM we avoid the unnecessary burden of coarse-graining a fine grained MSM whose microstates are certainly not fulfilling the assumption of Markovianity.

### B. Alanine dipeptide

Our second example is the terminally-blocked alanine residue (see Fig. 3A), which is realistic in the sense of being a molecular model that experiences slow transitions between conformational states, while still keeping a very small number of degrees of freedom. Specifically, here we use the *ϕ* and *ψ* Ramachandran angles. Again, we compare the results from the standard procedure for generating MSMs against our simplified hdbMSM workflow.

**FIG. 3:**
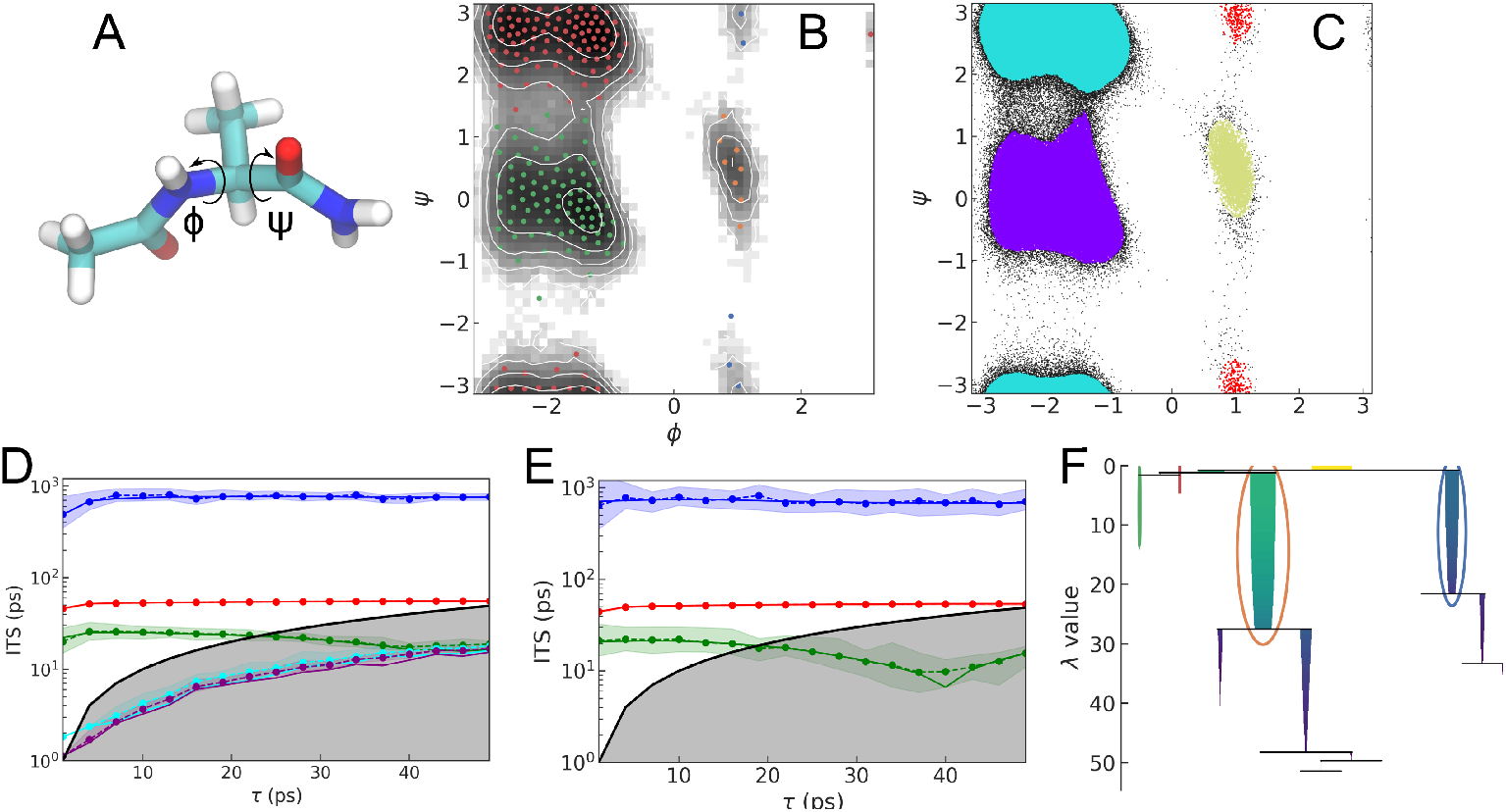
(A) Representation of the alanine dipeptide with the *ϕ* and *ψ* torsion angles. (B) Projection of the microstates of the fine-grained MSM on the Ramachandran map, overlaid on a density plot calculated from our simulation trajectories. Clusters were generated using *k*-means clustering with *k*=200. Colour code indicates the different metastable states obtained by using PCCA+ coarse-graining. (C) Data points from the simulation color-coded by cluster index as determined by HDBSCAN. Noise points are shown in black. We used a minimum cluster size of 350. (D) Implied timescales as a function of the lag time (both in ps) from the fine grained MSM. Solid lines are maximum likelihood estimates and dashed lines are mean estimates from Bayesian sampling. Shaded regions mark 95% confidence intervals. (E) Same for the hdbMSM. (F) Condensed tree plot and selected clusters obtained for alanine dipeptide by using HDBSCAN.

In Figure 3B we show the projection of the fine-grained MSM microstates on the Ramachandran map. As in the case of the bistable potential, the system has only two degrees of freedom, and hence we do not need to use dimensionality reduction techniques. In order to avoid problems from the periodicity of the backbone torsions, before the clustering we shifted the values of *ϕ >* 2 and *ψ* < – 2 by –2*π* and 2*π*, respectively. Then we clustered the data using the *k*-means method with *k* = 200 cluster centres. Using these clusters we obtain a fine grained MSM with three slow modes, which have a slight lag time dependence for *τ* < 5 ps, plus many other fast modes. Subsequently, we performed a PCCA+ coarse-graining, which results in four macrostates that match the dominant *α*-helix, PPII/*β* and *α*_L_ basins (see colour code of microstates in Fig. 3B).

When we directly cluster the values of *ϕ* and *ψ* using HDB-SCAN with a minimum cluster size of 350 we directly recapitulate the relevant free energy basins, leaving out as noise datapoints in boundary regions between the free energy minima (see Figure 3C and Fig. S4 in the Supplementary Material for the time series of the backbone torsion angles assigned to the different states). In order to be more stringent with the TBA procedure, we left as noise points those that belong to their clusters with probabilities lower than 0.3. Using this state assignment onto four unique states, we calculate the hdbMSM and determine the corresponding implied timescales, which are shown in Figure 3E. The resulting values are in quantitative agreement with the fine-grained MSM results (Fig. 3D) and are converged at the minimum lag time of 1 ps. The agreement between both workflows is outstanding, as both are able to capture the relevant timescales. However, the simplified procedure that we argue for is advantageous in that it skips the need of deriving the fine grained MSM where some of the microstates very much populate barrier regions between metastable basins (see Fig. 3B). We have performed Chapman-Kolgomorov tests to validate the models, and they are satisfactory both for the fine grained MSM and the hdbMSM (see Supplementary Information, Figs. S5 and S6).

While the value of the minimum cluster size of 350 captures a partitioning that results in a quantitatively accurate MSM, the choice of this parameter in HDBSCAN may not be straightforward in this example. In order to select its value, one can resort to the use of cluster tree plots^23,24,46^. According to the aforementioned description, condensed tree plots are obtained from the cluster hierarchy by saving only those clusters that are larger than the *minimum cluster size* parameter. As shown in Figure 3F and according to the explanation given in Sec. III B, the width of each cluster represents the number of points in the cluster and gets narrower as *λ* increases. Then, the clusters with largest persistence, which corresponds to a larger stability, are selected. In practice, the condensed tree plot must look interpretable, so that selected clusters have considerable width and there is no much noise at large *λ* values. In this particular example, two large clusters corresponding to the *α*-helix and PPII/*β* regions are dominant. Still, the two marginally populated free energy basins (i.e. *α*_L_ and *γ* basins) are also captured as shown in the top-left part of the tree in Figure 3F. This illustrates the ability of HDBSCAN to capture clusters even if they have drastically different shapes and densities.

### C. FiP35 WW domain

Finally, we apply the proposed workflow to the long timescale simulation trajectories (100 *µ*s each) of the FiP35 WW domain from the Shaw group^45^, a dataset used by others before in MSM development^47–49^. In this case, we first used dimensionality reduction with TICA^26–28^ using the distances between pairs of *α*-carbons as features. We keep three TICA dimensions, so that resulting components show significant transitions along the two independent trajectories (see Supplementary Material, Fig. S7). Following the standard procedure for building MSMs, we discretize the first few co-ordinates from TICA using *k*-means clustering with *k* = 100 cluster centers. The projection of the *k*-means clusters is shown on Figure 4A. The fine-grained MSM yields two slow implied timescales that are well converged only after, approximately, *τ* = 200 ns (see Fig. 4D). The implied timescales for these slowest modes from the fine grained MSM are approximately 5.5 and 1.5*µ*s, which compare well with previous results^47,48^. The MSM does not resolve faster timescales, as they are not converged for any *τ* < *t*_*i*_. Considering the resulting separation of time-scales, we use PCCA+ to coarse-grain the model into three macrostates (see Figure 4A).

**FIG. 4:**
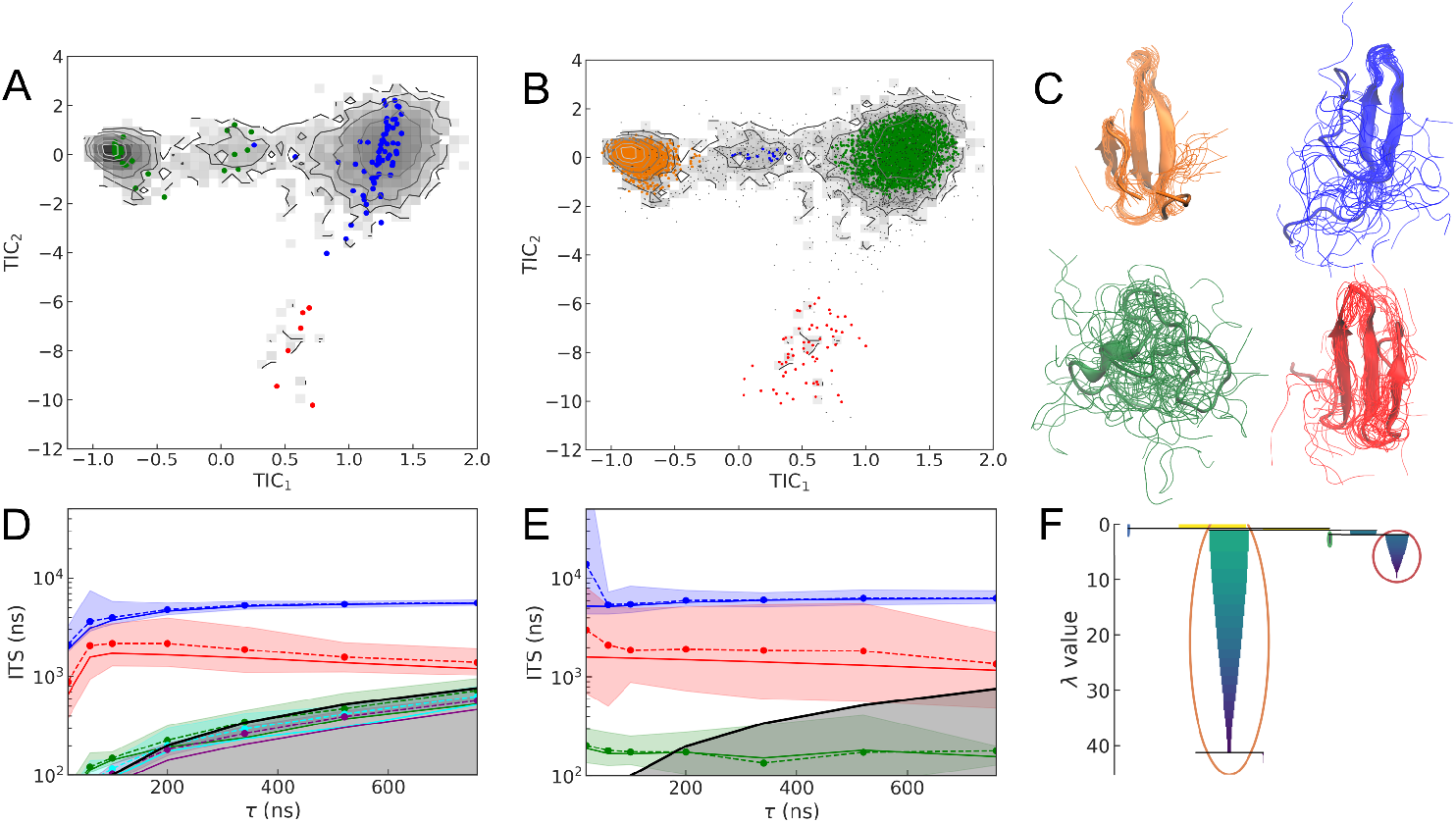
(A) Projection of the microstates of the fine-grained MSM of the FiP35 WW domain, overlaid on a density plot calculated from the long equilibrium trajectories. Clusters were generated using *k*-means clustering with *k*=100. Colour code indicates the different metastable states obtained by using PCCA+ coarse-graining. (D) Implied timescales as a function of the lag time from the fine grained MSM. Solid lines are maximum likelihood estimates and dashed lines are mean estimates from Bayesian sampling. Shaded regions mark 95% confidence intervals. (D) Same for the hdbMSM. (E) Cartoon representation of the clusters identified using HDBSCAN, with colours relative to panel B. Each state is represented by 20 randomly selected conformations. (F) Condensed tree plot for the hdbMSM of Fip35 WW domain.

In the case of the hdbMSM, we directly assign the trajectory to the four, instead of three, distinct states resulting from clustering with HDBSCAN (see Figure 4B). Hence, using this approach we are able to identify one more coarse state than with the PCCA+ coarse-graining. In Figure 4E we show representative snapshots for each of these states. It is noteworthy that the clusters captured by HDBSCAN have drastically different densities, as illustrated in the condensed tree plot (Figure 4F). We report the timescales for the hdbMSM in Figure 4D which are converged earlier than those from the fine-grained MSM (i.e. at *τ* ≃50 ns). This suggests that the improved state definition from the hierarchical clustering may result in an improvement with respect to the usual workflow. In fact, the hdbMSM is able to resolve a third slow mode that we have not obtained with the typical protocol. From the eigenvectors corresponding to the slowest three eigenmodes (see Supplementary Information, Fig. S10), we find that the additional mode corresponds to the exchange between the intermediate state shown in blue in Figure 4E, with two *β*-sheets formed, with the unfolded state and the fully folded state (green and orange, respectively). Corresponding CK tests are shown in Figures S8 and S9 in the Supplementary Information for the MSM and the hdbMSM, respectively.

## V. CONCLUSIONS

Here we show that, using a hierarchical density-based clustering method, HDBSCAN, we can make the workflow for MSM construction simpler than with the more established protocols^9^, while preserving the accuracy in the estimation of the slow dynamics. In previous applications of density-based approaches to MSM construction, DBSCAN was shown to be well-suited for the construction of core-sets MSMs. However, in these applications the density-based method simply replaced the popular, centroid-based *k*-means clustering method. In this work, we aim to go a step beyond and employ HDBSCAN to directly coarse-grain the feature space either from trivial coordinates or from optimal coordinates resulting from dimensionality reduction in complex molecular systems. Thus, we avoid defining microstates and building a fine-grained MSM, which in any case rests on the false assumption of Markovian dynamics. Additionally, we do not require a subsequent coarse-graining step using PCCA+ or similar methods for lumping microstates. In other words, HDB-SCAN directly produces an accurate and interpretable model with only a few states with high metastability. The resulting protocol for building MSMs is referred to as hdbMSM.

We choose three application examples of increasing complexity to validate our methodology. By studying a toy model and alanine dipeptide we demonstrate, respectively, the ability to identify metastable regions and to describe the slow kinetics among well-defined core-sets. For these examples, the standard procedure for MSM construction also works well, but we are able to reproduce the same result using a more simplified workflow. On the other hand, a well-established complex biological system serves as a challenging problem for the approach that we propose. For the FiP35 WW domain the hdbMSM is converged for shorter lag times than in the conventional approach. More importantly, the third slow mode is only retrieved by hdbMSM, whereas the standard procedure for building MSMs cannot capture it. In fact, this transition requires to identify a cluster of low density associated with two *β* strands which only HDBSCAN is able to account for.

We do not attempt to introduce a full automation of the hdbMSM workflow, as recently done in the context of deep-learning approaches. However, our approach may proof helpful for current MSM analyses of MD data with openly available tools as it simplifies the construction of MSMs, reduces the input parameters from the end user and enables shorter lag times. It must be emphasised that our work is necessarily limited to a few application examples. All the systems considered have significant gaps among different modes. It remains an open question if hdbMSM, and more generally density-based clustering algorithms, will yield accurate results for systems with a more continuous spectrum of eigenmodes.

## DATA AVAILABILITY

The materials that support the findings of this study are openly available at https://github.com/BioKT/hdbMSM. Simulation trajectories corresponding to alanine dipeptide are available from the corresponding author upon request.

## ACKNOWLEDGMENTS

We acknowledge D. E. Shaw Research for sharing the long timescale MD trajectories of FiP35 WW. Financial support to DDS comes from Eusko Jaurlaritza (Basque Government) through the project IT1254-19, Grants RYC-2016-19590 and PGC2018-099321-B-I00 from the Spanish Ministry of Science and Universities through the Office of Science Research (MINECO/FEDER) and the Dosostia International Physics Center (DIPC). We also acknowledge the staff at the DIPC Supercomputing Center for technical support.

