## Supporting Information for "Markov state models from hierarchical density-based assignment"

### Supplementary material for
### Markov state models from hierarchical density based assignment

Ion Mitxelena,[1] Xabier López,[1] and David de Sancho[1, a)]

*Polimero eta Material Aurreratuak: Fisika, Kimika eta Teknologia, Kimika Fakultatea, UPV/EHU & Donostia International Physics Center (DIPC), PK 1072, 20018 Donostia-San Sebastian, Spain*

(Dated: 13 May 2021)

[a)]Electronic mail:

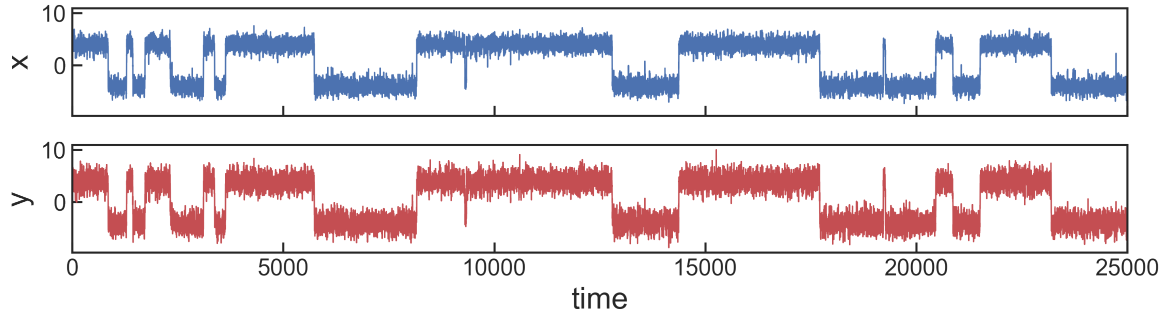


FIG. S1: Time series data corresponding to the brownian dynamics trajectory on a two-dimensional potential.

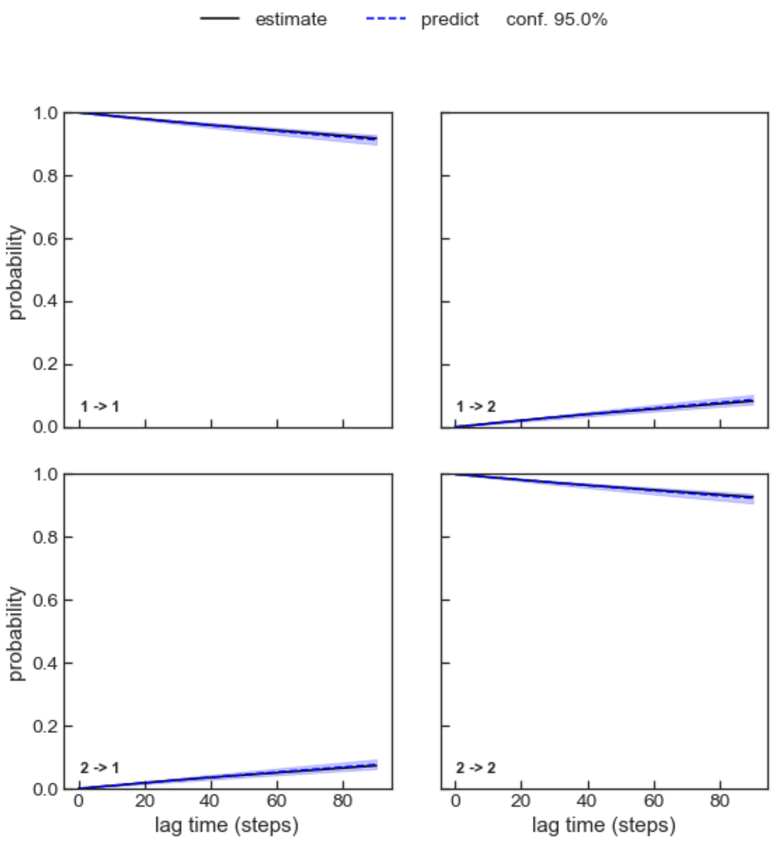


FIG. S2: Chapman-Kolmogorov tests corresponding to the two-dimensional potential from PCCA+.

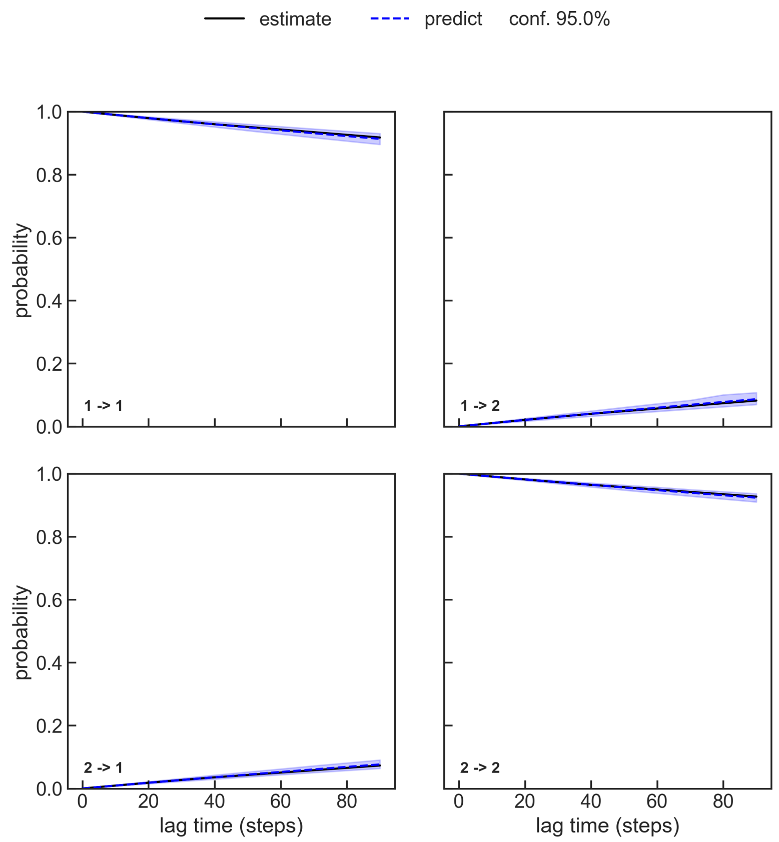


FIG. S3: Chapman-Kolmogorov tests corresponding to the two-dimensional potential for the hdbMSM.

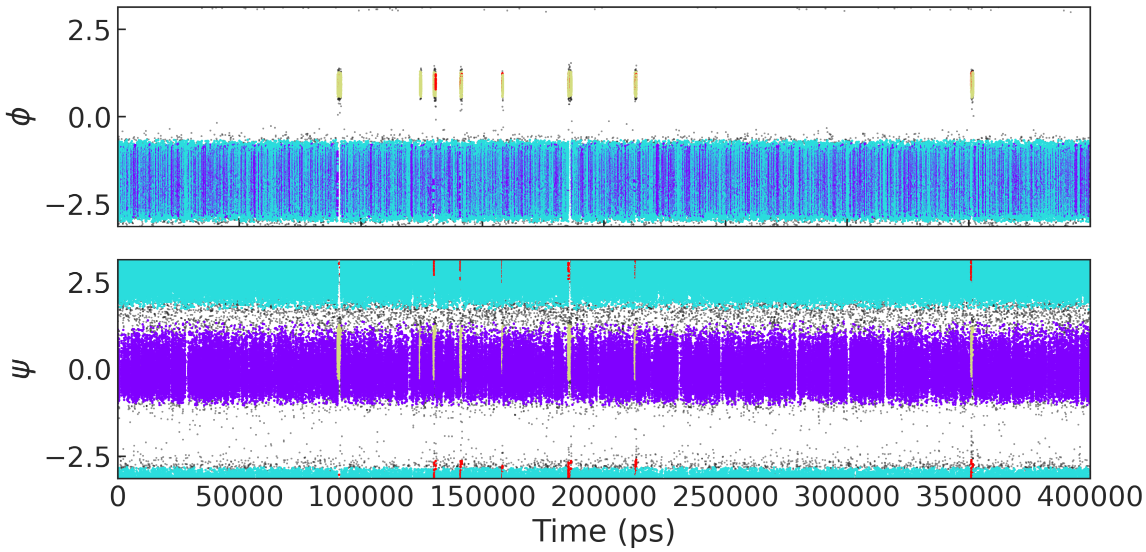


FIG. S4: Backbone torsion angles with different colours for each HDBSCAN cluster along simulation trajectory of alanine dipeptide.

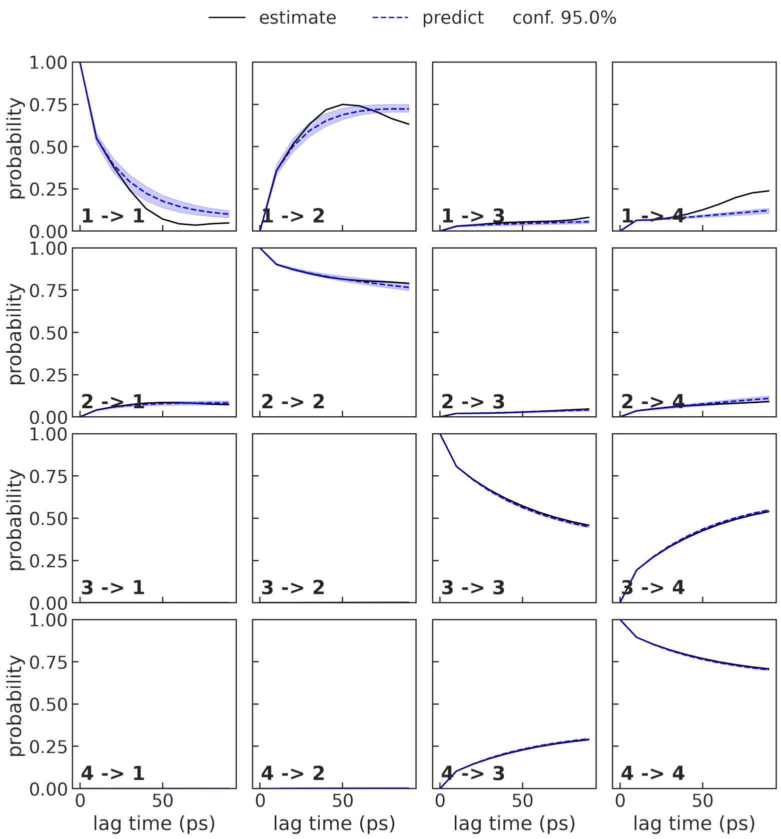


FIG. S5: Chapman-Kolmogorov tests corresponding to alanine dipeptide. for the PCCA+ coarse-grained MSM.

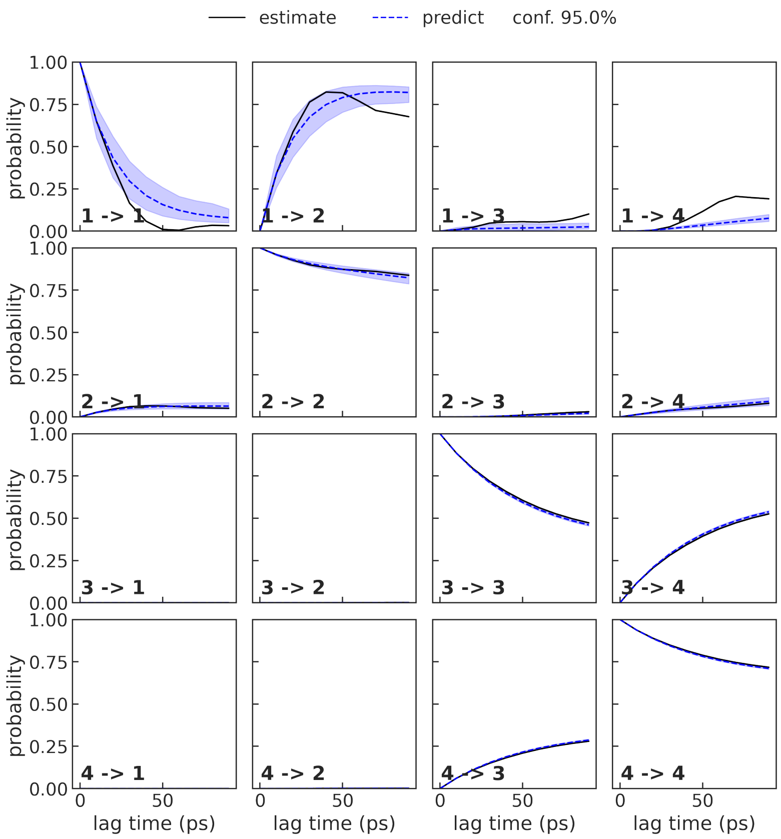


FIG. S6: Chapman-Kolmogorov tests corresponding to the hdbMSM for the alanine dipeptide

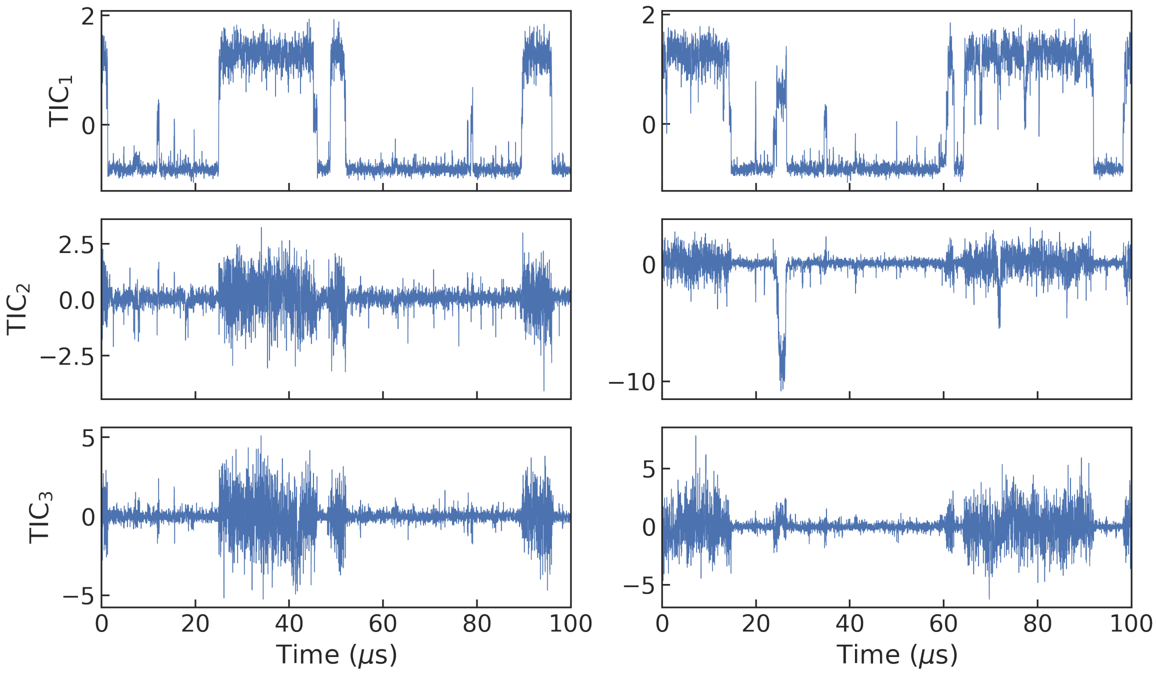


FIG. S7: Time series data for the projection on the first three TICA coordinates for the two long equilibrium trajectories of the FiP35 Pin WW domain.

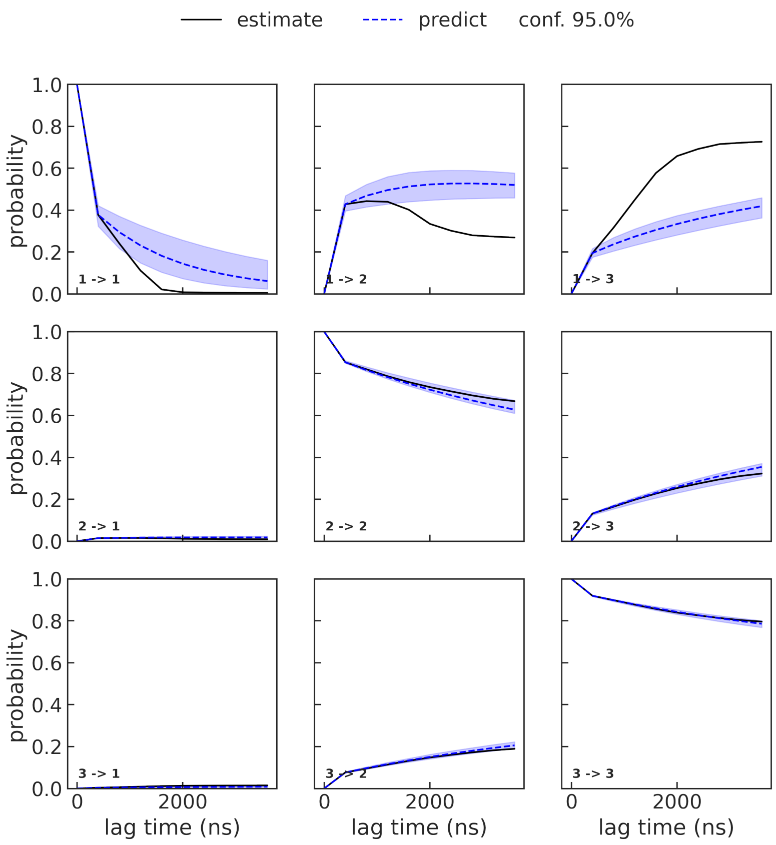


FIG. S8: Chapman-Kolmogorov tests corresponding to the FiP35 WW domain from the MSM coarse-grained using PCCA+.

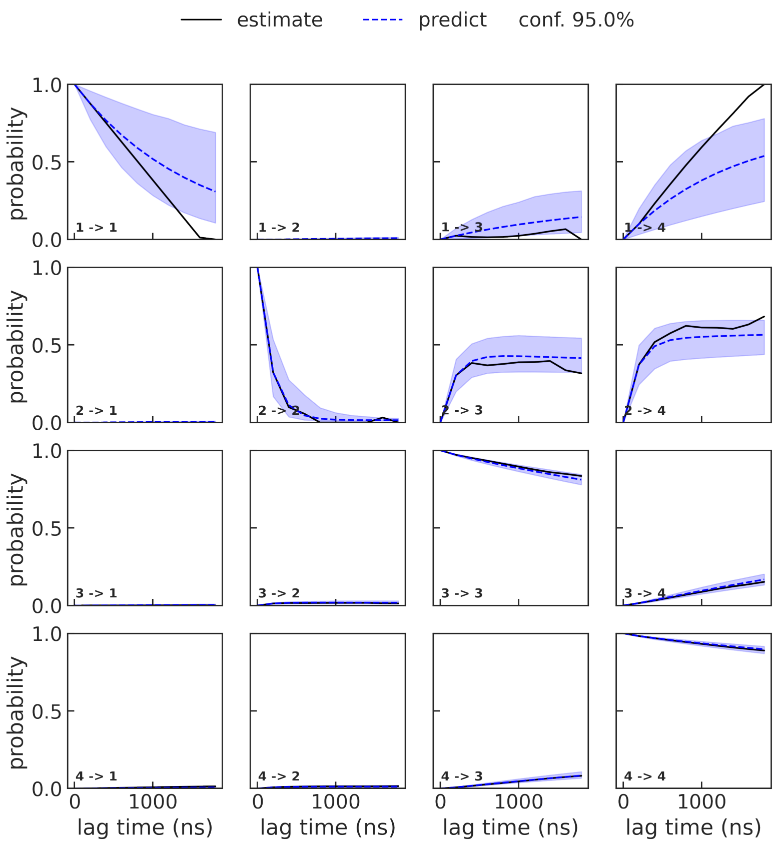


FIG. S9: Chapman-Kolmogorov tests corresponding to the hdbMSM for the FiP35 WW domain.

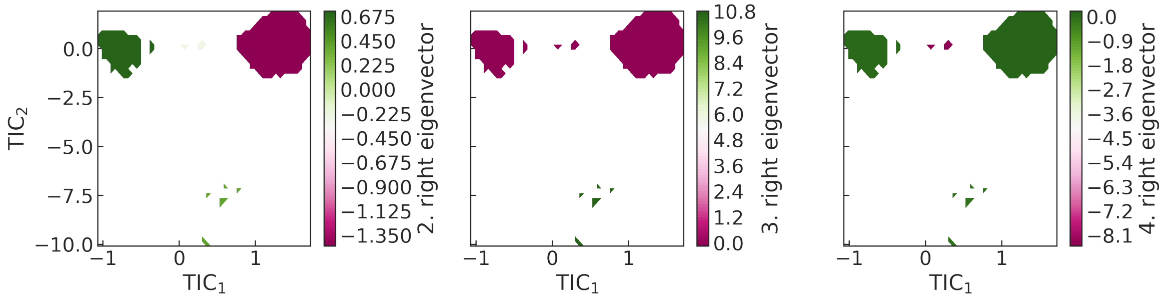


FIG. S10: Right eigenvectors corresponding to the three slowest modes of the hdbMSM for the Fip35 WW domain.
